# Pronoun resolution via reinstatement of referent-related activity in the delta band

**DOI:** 10.1101/2023.04.16.537082

**Authors:** Rong Ding, Sanne ten Oever, Andrea E. Martin

**Affiliations:** Language and Computation in Neural Systems, Max Planck Institute for Psycholinguistics, Nijmegen, The Netherlands; Donders Centre for Cognitive Neuroimaging, Nijmegen, The Netherlands; Department of Cognitive Neuroscience, Faculty of Psychology and Neuroscience, Maastricht University, Maastricht, The Netherlands

**Keywords:** pronoun resolution, reinstatement, memory, neural oscillations, delta band, spoken language comprehension, MEG

## Abstract

Human language offers a variety of ways to create meaning, one of which is referring to entities, objects, or events in the world. One such meaning maker is understanding to whom or to what a pronoun in a discourse refers to. To understand a pronoun, the brain must access matching entities or concepts that have been encoded in memory from previous linguistic context. Models of language processing propose that internally stored linguistic concepts, accessed via exogenous cues such as phonological input of a word, are represented as (a)synchronous activities across a population of neurons active at specific frequency bands. Converging evidence suggests that delta band activity (1-3Hz) is involved in temporal and representational integration during sentence processing. Moreover, recent advances in the neurobiology of memory suggest that recollection engages reinstatement of neural dynamics that occurred during memory encoding. Integrating from these two research lines, we here predicted that neural dynamic patterns, especially in delta frequency range, underlying referential meaning representation would be reinstated during pronoun resolution. By leveraging neural decoding techniques (i.e., representation similarity analysis) on a magnetoencephalogram (MEG) dataset acquired during a naturalistic story-listening task, we provide evidence that delta-band activity underlies referential meaning representation. Our findings suggest that, during spoken language comprehension, endogenous linguistic representations such as referential concepts may be retrieved and represented via reinstatement of dynamic neural patterns.

## Introduction

Consider the meaning of the pronouns *she* and *herself* in the following passage:

> “The **fool** doth think [she] is **wise**, but the wise [woman] knows [herself] to be a fool.”
>
> — Adapted from Shakespeare’s *Twelfth Night*

To understand the above passage, we have to combine the previously presented story agent (i.e., “the fool”) with the pronoun *she* to understand that it is the fool who considers herself wise; then we must understand that *herself* refers to *the wise woman* but not *the fool*. Thus, to comprehend the sentence, or any event involving a pronoun, some form of memory of the matching antecedent must come into play when the pronoun is processed. Pronouns are ubiquitous in human language and their use and interpretation is a cornerstone of human cognition (e.g., Garrod & Sanford, 1994) and development (e.g., Hendriks & Spenader, 2006). But how this quintessential linguistic device is realized in neural terms, such that it enables us refer to agents, events, and concepts that span time and space, is not at all well understood. Furthermore, pronoun processing sits at the intersection between language and memory, offering a fruitful way to study the interplay between these two types of information in the brain.

A burgeoning cue-based account of language in the brain (e.g., Martin, 2016, 2020) has proposed that comprehension is a perceptual inference process whereby the brain takes exogenous cues (e.g., sensory features) as its input and combines them with endogenously generated linguistic cues (e.g., lexical entries, procedural syntactic rules) from memory to achieve comprehension. Pronoun resolution thus also fits into this account as it requires access to previously encoded entities or concepts in order to integrate them in turn with roles the pronoun plays, so that coherent construction of events in a text or conversation can be achieved. Indeed, supporting evidence has emerged from a plethora of studies showing that properties of both external and internal cues (e.g., featural match/mismatch, referent prominence), as well as how they are combined, can influence how the brain resolves a pronoun (LeDoux et al., 2007; Foraker & McElree, 2007; Parker, 2019; Nieuwland & Van Berkum, 2008; Nieuwland, 2014; Chow, Lewis, & Philips, 2014; Brodbeck, Gwilliam, & Pylkkanen, 2016; Brodbeck & Pylkkanen, 2017; Karimi, Swaab, & Ferreira, 2018; Lissón et al., 2021; Nieuwland, Coopmans, & Sommers, 2019; Coopmans & Nieuwland, 2020). Yet, the neural mechanism by which a referent concept is retrieved and represented in memory when the brain resolves a pronoun has barely been discussed. Major neurobiological findings regarding pronoun resolution come from studies using the event-related potential (ERP) technique, which have identified an increased, sustained anterior negative component (i.e., Nref) induced in pronouns which are incongruous with their antecedent (e.g., Nieuwland, 2014) or ambiguous in their reference (e.g., Nieuwland & Van Berkum, 2008; Karimi, Swaab, & Ferreira, 2018; Coopmans & Nieuwland, 2020). However, while these results provide evidence in favor of the cue-based account by showing that retrieving internally stored entity representations can be interfered by processing referentially incoherent or ambiguous input, they do not give a clear picture how the brain actually accesses referent concepts. Besides, the modulation of neural activities by mismatch or ambiguity of pronouns are subject to other possible interpretations based on domain-general cognitive functions, for instance enhanced attentional process due to error or ambiguity detection. In other words, a converging mechanistic account of how the retrieval of a referent concept from memory is accomplished in the brain remains a missing puzzle from existing studies on referential resolution.

Recent advances in the domain of neuroscience of memory have provided insight into memory retrieval through neural decoding techniques. Numerous findings have indicated that neural patterns during memory encoding are reinstated during memory retrieval (e.g., Rugg & Johnson, 2007; Manning et al., 2012; Yaffe et al., 2014; Jang et al., 2017; Xiao et al., 2017; Yaffe et al., 2017; Pacheco Estefan et al., 2019; Staresina et al., 2019; Ten Oever et al., 2021). In particular, oscillatory neural dynamics, often referred to as putative neural oscillations and postulated to arise from an ensemble of neurons firing synchronously, have been found reinstated during recollection (e.g., Nyhus & Curran, 2010; Yaffe et al., 2014; Yaffe et al., 2017; Staresina et al., 2019; Ten Oever et al., 2021). Among these studies, representational similarity analysis (RSA), i.e., a measurement of the similarity between neural states during memory encoding and retrieval acquired through multi-variate pattern analysis, has been applied to investigate the reinstatement of previous activity patterns as a brain mechanism for retrieval. Relevant to language processing, such effects have also been observed during word retrieval (Yaffe et al., 2014; Yaffe et al., 2017; Ten Oever et al., 2021). It has been shown that theta-(3.5-8Hz) and gamma-band (50-100Hz) activity underlying word learning was reinstated during successful word recall in a verbal association task (e.g., Yaffe et al., 2014; Yaffe et al., 2017). Additionally, in this same line of research, converging evidence has shown that the temporal lobes, including both lateral temporal (e.g., the fusiform gyrus, middle temporal gyrus; Rugg & Johnson, 2007; Jang et al., 2017) and medial temporal regions (e.g., hippocampus; Manning et al., 2012), are engaged in reinstatement of memory traces (Pacheco Estefan et al., 2019; Staresina et al., 2019) and, importantly, lexical retrieval (Ten Oever et al., 2021). Altogether, these results provide evidence that the reinstatement of neural-oscillatory responses underpinning memory traces in both lower- and higher-frequency bands (e.g., theta and gamma), particularly across temporal regions, is involved as memory representations during retrieval.

Consistent with memory findings indicating that oscillatory dynamics play a role in memory retrieval, the cue-based retrieval account of language processing purports that linguistic cues, either external or internal, can be represented by neural oscillations (e.g., Martin, 2016, 2020) and serve to elicit or “serve up” existing information in the brain – implicitly, this claim indicates that that process must rely on some forms of memory. Indeed, a body of research bridging neural oscillations and language processing has suggested that the human brain’s ability to form representations from abstract linguistic symbols is enabled by multiplexing of its rhythmic activities on various timescales to impose internal knowledge upon external, incoming input (Kösem & van Wassenhove, 2016; Meyer, 2018; Rimmele et al., 2018; Meyer, Sun, & Martin, 2019; Gwilliams, 2020; Martin, 2020; Martin & Doumas, 2017, 2019, 2020). It has been shown that neural oscillations are involved in not only the processing of sensory streams (e.g., Luo & Poeppel., 2007; Kösem et al., 2018) but also in the processing of higher-order linguistic representations (e.g., Bai, Meyer, & Martin, 2022; Brennan & Martin, 2019; Coopmans et al., 2022; Ding et al., 2016; Meyer et al., 2017; Kaufeld et al., 2020; Henk & Meyer, 2021). Concretely, delta-band activity (1-3Hz) has been found to be relevant for the tracking of meaningful linguistic elements such as words and phrases (e.g., Ding et al., 2016; Meyer et al., 2017; Kaufeld et al., 2020; Henk & Meyer, 2021; Ten Oever et al., 2022). However, it remains an open question how activity in the delta band is involved in this process, or rather, it remains unclear what precisely modulations of delta represent in terms of neural information processing. Besides the role of tracking higher-level linguistic components from sensory input as suggested by the aforementioned studies (also see Rimmele et al., 2018; Lakatos, Gross, & Thut, 2019), another postulated role of delta has been proposed, i.e., as a functional pattern that reflects the generation of abstract linguistic representations (e.g., Martin, 2020; Meyer, Sun, & Martin, 2019; Ten Oever & Martin, 2021). This creation or generation of structure and information likely entails recognition, reactivation, or retrieval of information from memory. In the current study, by investigating the neural-oscillatory substrates of pronoun resolution we aim to focus on this postulated, top-down function of delta in imposing previously encoded memory representations on incoming speech. Pronoun processing necessarily requires the brain to recover higher-order linguistic elements that are previously stored in memory, instead of simply tracking them directly from the sensory input of a pronoun *per se*. In this case, if modulations of delta-band activity, importantly reinstatement of delta activity underlying referent representation were indeed found when the brain resolves a pronoun and constructs the corresponding event, then it would support the proposal that delta activity is involved in the retrieval of stored higher-order linguistic representations or in the integration of that information with its current role in the sentence or discourse. Given that pronouns can vary in the time duration they take up in speech processing, both within and across languages, finding evidence for effects of pronoun retrieval and integration in the delta band would support a functional account of frequency bands that is not necessarily tied to the absolute, external timing of stimulus presentation, but rather more endogenous and abstract, during language comprehension. Therefore, investigating the rhythmic neural responses underlying pronoun resolution serves as a promising opportunity to better understand the mechanistic roles oscillatory neural dynamics play in language processing.

Therefore, in light of oscillatory neural activation reinstatement as an emerging account of memory retrieval, the current study seeks to investigate whether referent concept representation during pronoun resolution also engages the reinstatement of oscillatory activities underlying antecedent processing in the brain. Though separately, existing findings have suggested the involvement of neural pattern reinstatement and oscillatory responses in pronoun resolution (Zhang et al., 2022; Nieuwland & Martin, 2017; Coopmans & Nieuwland, 2020). Several EEG findings (Nieuwland & Martin, 2017; Coopmans & Nieuwland, 2020) identify enhanced power of rhythmic brain responses such as gamma when the brain processes pronouns coherent with their corresponding antecedent. Meanwhile, a decoding study by Zhang et al. (2022) found activation in the left MTG that underpinned presentation of story characters during processing of zero pronouns (i.e., an obligatory noun phrase that serves a role in an event but is not overtly pronounced in the utterance) in Mandarin Chinese. By leveraging RSA, the study managed to zoom in and compare neural responses between individual items (i.e., each referent and zero pronoun), which have largely been smeared out by grand-averaged patterns of two coarse lexical categories as in traditional condition-based statistical analyses. This way, hypotheses about item-specific neural fluctuations (in this case reinstatement) were tested. Therefore, by adopting RSA as the neural decoding technique, the current study aimed to provide first direct evidence that oscillatory neural responses underlie referent representation via reinstatement during pronoun processing.

The main question we ask in this study is whether rhythmic neural patterns underlying processing of the antecedent of a pronoun are reinstated when that pronoun is resolved during spoken language comprehension. To answer the question, we conducted RSA on responses to pronouns and their noun antecedents extracted from a magnetoencephalogram (MEG) dataset which was recorded while participants listened to continuous audiobook stories in Dutch. We predict that neural responses associated with higher-order linguistic elements, i.e., modulations of the delta frequency range – in this case, those responses associated with the antecedent noun – will recur systematically during pronoun resolution, and thus reinstating the rhythmic neural patterns of the antecedent during pronoun resolution. Besides, given that theta- and gamma-band activity have also been found associated with word retrieval (e.g., Yaffe et al., 2014; Yaffe et al., 2017; Ten Oever et al., 2021) and also referential resolution (e.g., Nieuwland and Martin, 2017; Coopmans & Nieuwland, 2020), we predict that reinstatement of theta and gamma responses that underlie referential noun processing will also be observed when the brain processes a pronoun.

## Method

### Participants

29 participants (21 females; mean age: 37.14 years old) with normal hearing and no history of psychiatric, neurological or other medical illness that might compromise cognitive functions took part in the experiment. They identified themselves as native Dutch speakers and self-reported having little or no prior knowledge of French. All participants gave informed consent before the experiment and received monetary compensation for their participation. This study was approved by the ethical Commission for human research Arnhem/Nijmegen (project number CMO2014/288). Two participants did not complete the experiment, and one other wore dental wires during the recording session. Datasets of the three participants were thus removed from further analyses.

### Stimuli and Paradigm

In the experiment, participants were instructed to listen to audiobook stories while their MEG responses were being measured. The audiobook stimuli consist of 3 Dutch stories and 3 French stories presented in a pseudo-randomised order. The Dutch stories include Anderson’s *Het Leelijke Jonge Eendje*, and Grimm’s *De Ransel, het Hoedje en het Hoorntje* and *De gouden vogel*, and the French stories Anderson’s *L’Ange*, Grimm’s *L’eau de La Vie*, and E.A. Poe’s *Le Canard au ballon*. In this study we focus on the Dutch stories. The French stories were collected as part of a long-term commitment to collect a bigger MEG set of naturalistic audiobook listening including the option for cross-language comparison which was not part of this study.

All stories were split into blocks lasting approximately 5 minutes each; this way, participants were each presented with a total of 13 blocks (9 blocks for Dutch stories and 4 for French stories). Prior to each block, each participant’s resting state brain activity was recorded for 10 seconds. Between each two blocks participants were indicated to answer 5 multiple-choice comprehension questions based on the story content they have just heard to ensure they paid attention to the stimuli. Including set-up and breaks, the entire MEG session took an average of 90 minutes.

Prior to the experiment, each participant completed a 5-minute, MEG auditory localiser task whereby they listened to tones while the brain responses were recorded. They also underwent MRI structural scanning before or after the MEG session.

### Data recording

MEG data were recorded using a 275-channel whole-brain axial gradiometer system (CTF VSM MedTech) at a sampling rate of 1200 Hz. Bipolar vertical and horizontal EOG and ECG were recorded using Ag/AgCl-electrodes. Six channels were permanently faulty and two others successively disabled during the recordings, leaving 269 recorded MEG channels for 23 participants and 267 for three participants. Head localization was monitored continuously during the experiment using fiducial coils that were placed at the cardinal points of the head (nasion and bilateral ear canals). The fiducial coils also served as anatomical landmarks for co-registration with MRI scans during source reconstruction. Stimulus presentation was controlled by the Matlab toolbox *Psychtoolbox*. Immediately after the MEG session, each participant’s headshape and position of three fiducial points were recorded using a Polhemus 3D tracking device (Polhemus, Colchester, Vermont, United States).

The T1-weighted MRI scan of each participant was acquired on one of three 3T MRI scanners (Siemens) available at the Donders Institute. Earplugs with a drop of vitamin E were placed at the subject’s ear canals during MRI acquisition, to facilitate co-registration between MEG and MRI data.

### Analyses

#### Target word Selection

For pronoun word stimuli, we selected all Dutch personal, possessive, and reflexive pronouns (except second-person ones). For referent words, we picked out the main noun of noun phrases referred to by the chosen pronoun words. In the end, 791 pronouns and 407 referent nouns in the Dutch audiobook stories were selected as target words.

#### Preprocessing and time-frequency analyses

MEG data were preprocessed using Fieldtrip toolbox (Oostenveld et al., 2011) running under Matlab 2021a (The Mathworks, Inc.). Two participants were excluded from further analyses due to incompletion of story listening, and one other due to noise caused by metal magnetisation identified in recorded data. Data were first epoched as -2 to 3.5s around word onset and downsampled to 300Hz with an anti-aliasing low-pass filter applied prior to resampling. Artefact detection and rejection was conducted to remove trials that contained muscular and jump noise; during this process the artefacts were first identified automatically and then subjected to manual rejection. Independent component analysis (ICA) was then performed to remove components including ocular movements, slow drifts, heart beat and other salient noise. All the datasets were then subjected to a time-frequency analysis, yielding both power and phase. Activity at low frequencies (1-30Hz) underwent a wavelet analysis (1-Hz step size), with variable widths from 3 to 10 cycles (linearly increasing across frequencies) and a time range of -0.5 to 2 seconds (0.05-s step size). For higher frequencies (35-150Hz), a multitaper analysis (dpss taper; 10 cycles data at a 0.5-cycle smoothing) was conducted with a step size of 5Hz. All power values from the analyses were log-transformed by single trial per frequency. We chose to not conduct baseline correction as 502 of the 1198 trials have at least one target word stimulus from other trials present within 1.5 seconds pre-word onset. Baseline correction therefore would possibly introduce biases in the observed difference between the two conditions.

#### Sliding-window RSAs

In light of previous findings suggesting that the temporal cortex is involved in reinstatement and maintenance of lexical representations (e.g., Jang et al., 2017; Ten Oever et al., 2021; Zhang et al., 2022; Yaffe et al., 2017), in the current study we used bilateral temporal channels. Spearman correlations were computed for each pair of referent and pronoun trials across all sensor channels in the bilateral temporal regions. To investigate the specific time periods in which the effect takes place, we adopted a sliding-window approach on time axes of both referent and pronoun trials (see **Figure 1**); that is, for a pair of a pronoun and a referent trial, a 50-ms wide sliding window was moved on the two trials respectively, in a time range between 0 to 1.5 seconds. Then, correlations were computed between vectorised frequencies * channels * time datapoints of the two trials in a pair for each of the 50-ms time bins. This resulted in a total of 900 (30*30) pronoun x referent time*time units in a temporal generalization map. By contrasting between the values produced by correlating matching and non-matching pronoun-referent pairs, we tested the similarities between neural activities of referent and pronoun processing, i.e. the reinstatement of referent representations during pronoun resolution. The correlations and cluster-based statistics were conducted on the delta band (1-3Hz) as our band of interest. Spearman correlations were conducted on power values, and circular-circular correlations on phase values using the *CircStat* toolbox (Berens, 2009). Any pair of words whose onset difference was within 1.5 seconds was also excluded. Importantly, we selectively included non-matching referent-pronoun word pairs for the computation of similarities so that the median word count distances in the two conditions remained identical. This was to control for the similarities between neural activities due to adjacency in time (which can inflate correlations). Note that criteria were set only for non-matching referent-pronoun pairs because they were substantial in number, plus removing those of them in which the two words were distant from each other was already sufficient to keep the medians identical. Besides, as some trial epochs overlapped with each other in time, we excluded from statistical analyses the referent-pronoun trial pairs wherein other stimulus words were included that had the same reference. This ensured that the similarity effects we observed were only due to the processing of the word pairs that defined the trials.

**Figure 1.**
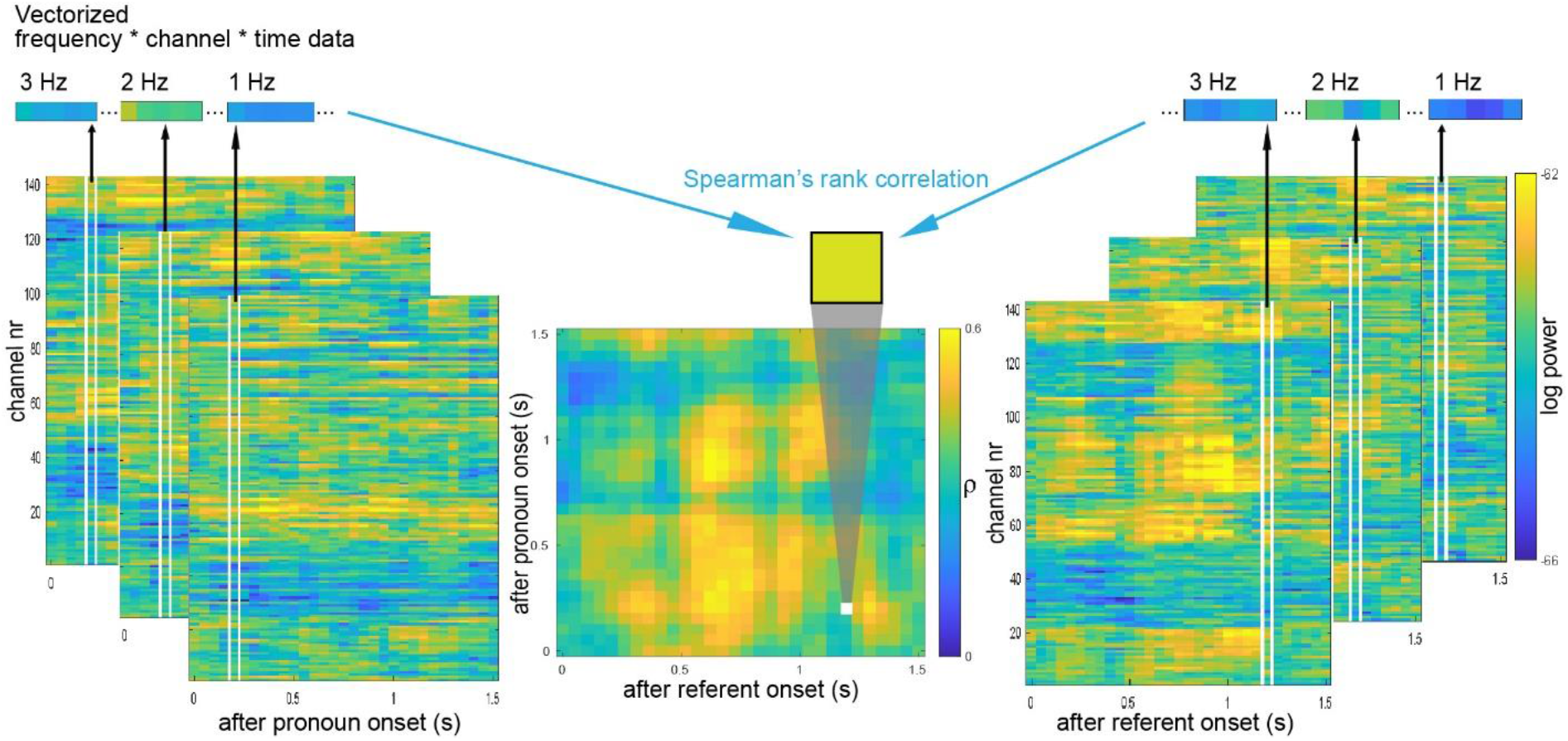
Computing reinstatement for each pair of pronoun and referent trials. The plots on the left and right show respectively time-frequency power patterns of the delta band (1-3Hz) of a pronoun and a referent trial. To compute the similarity between the two trials at a certain time*time unit, vectors of all frequencies (per band) * channels * time points were built. For power, the Spearman’s rank correlation coefficient between the pronoun and referent trial vectors determines the value of a single datapoint (i.e., time*time point) in a temporal generalization map. For phase, the value is determined by the circular-circular correlation coefficient, computed using *CircStat* toolbox (Berens, 2009).

Further, to better understand the temporal dynamics of reference resolution that were identified in pronoun-referent RSAs, we also conducted a control RSA between referent words and between pronoun words respectively (namely, word control RSAs). Concretely, we compared similarity values between pairs of words identical in form and those composed of differently formed words, irrespective of their reference. This should reveal the neural patterns evoked by perceptual and semantic features of single words. Same as on pronoun-referent stimulus word pairs, we conducted here the median word count control between identical and non-identical words pairs, as well as the exclusion of trial pairs where one trial contained overlapping word stimuli identical to any word stimulus in the other. As a control analysis, we repeated the analysis on theta (4-7Hz), alpha (8-12Hz), beta (13-20Hz) and low-gamma (35-80Hz) as well.

#### Statistical testing

Averaged matching and non-matching pronoun-referent (or word-word) correlation values were compared statistically using dependent-sample t-tests across participants in each pronoun x referent time*time unit. Cluster-based permutation tests were performed to correct for multiple comparisons (Maris & Oostenveld, 2007). Clusters were defined as a group of neighbouring time*time units with a p-value lower than 0.05 each in the dependent-sample t-tests. The sum of t-values of time*time units in a cluster was defined as the dependent variable on the cluster-level statistics. A distribution was then created by randomly permuting condition labels for 10000 times across participants and recomputing the test statistics produced by each permutation. The surrogate clusters with the maximum summed t-values entered the null distribution. Significance level (P-values) was then defined as the proportion of surrogate clusters in the distribution whose summed t-values were higher than those of the cluster observed from the actual data. We rejected null hypotheses when P-values were smaller than 0.05.

## Results

### 1 Delta band (1-3 Hz)

#### 1.1 Phase

In the pronoun-referent phase RSAs, a significant stronger Spearman correlation was found for the matching versus non-matching pronoun-referent pairs (Figure 2Ai; p = 0.0443). These results suggested that the phase of delta-band oscillations during referent representations was reinstated during pronoun resolution. Concretely, a reinstatement of delta phase between 350 and 700 milliseconds after referent onset took place in a time window of 750 to 1100ms post-pronoun onset.

**Figure 2.**
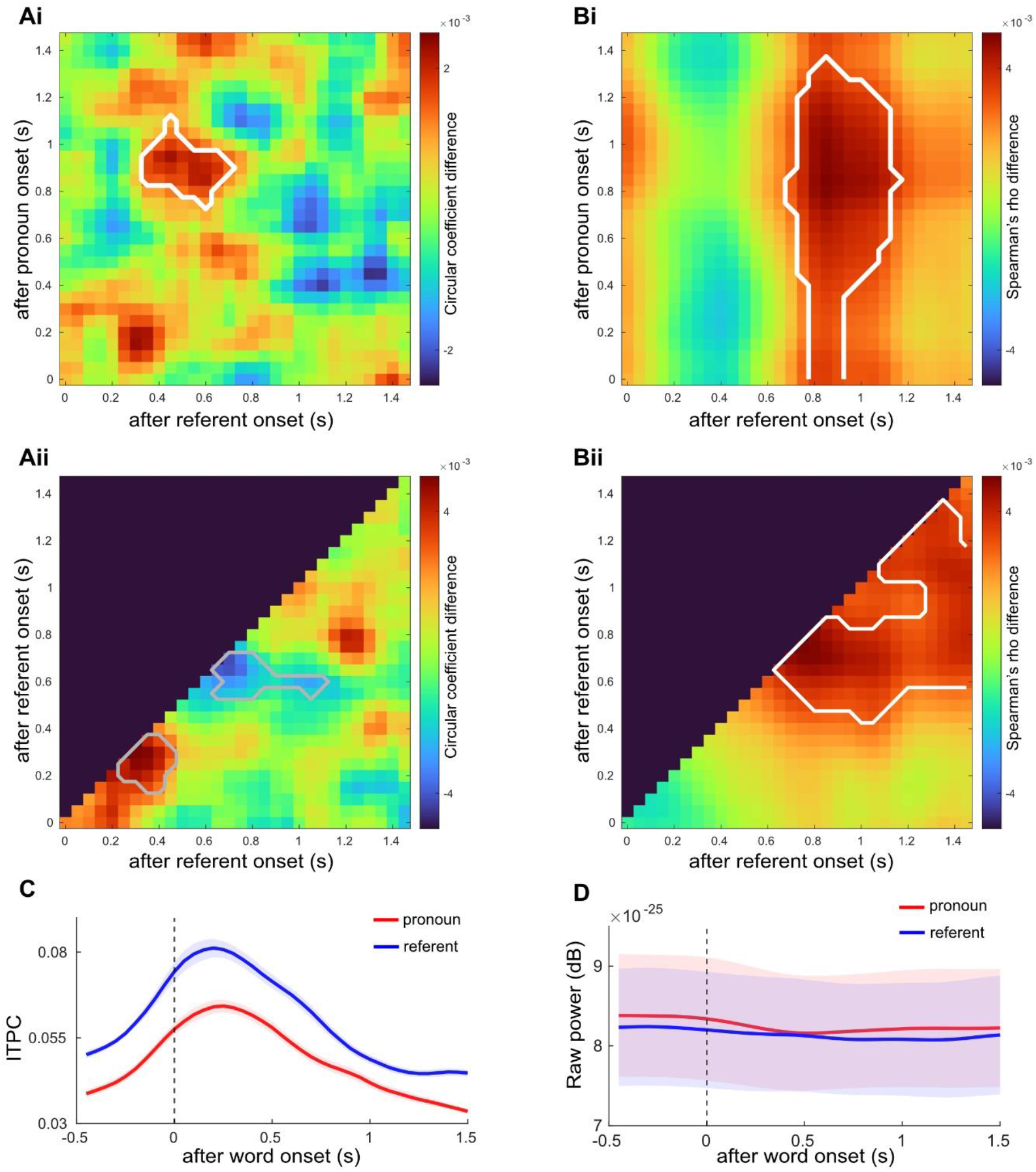
**Reinstatement of delta-band (1-3Hz) activities during pronoun resolution**. A) Reinstatement of neural representations of referent words is found in delta-band phase during pronoun resolution. (i) Temporal generalization map averaged across participants for the difference between matching and non-matching pronoun-referent word pair conditions. (ii) Averaged temporal generalization map for the difference between formally identical and non-identical word pair conditions. B) Reinstatement of neural representations of referent words is found in delta-band power during pronoun resolution. (i) Temporal generalization map averaged across participants for the difference between matching and non-matching pronoun-referent word pair conditions. (ii) Averaged temporal generalization map for the difference between formally identical and non-identical word pair conditions. C) Uncorrected, mean inter-trial phase coherence (ITPC) averaged per temporal channel per delta frequency. The ITPCs of pronoun and referent trials were plotted as separate lines. Note that the higher ITPC for referent words here could be due to the fact that the number of referent word trials are smaller than that of pronoun word trials. D) Mean delta power across the delta band and across all temporal channels. The mean power of pronoun and referent trials was plotted as separate lines. Shaded error bars indicate the standard error of the mean. Regions highlighted with white outlines indicate significant difference at the P = .05 level, while those highlighted in gray indicate trend-significant difference.

Word control RSAs between referent words yielded a trend-significance delta-band phase pattern activation between 200 and 450ms time-locked to referent word onset (Figure 2Aii; P = 0.0879). In addition, though, a trend-significance cluster where delta phase is more similar between non-matching referent and pronoun words than between matching pairs was also found (Figure 2Aii; P = 0.0522); the cluster went between 600 and 1100ms post-referent onset and between 550 and 750ms post-pronoun onset. No significant cluster was observed in the delta-band control RSA between pronoun words. A control analysis using source-level RSA, yielded no significant effect for neither the pronoun-referent or control comparisons (see Supplementary Materials for detail).

#### 1.2 Power

For the RSA between pronouns and referents, a significant cluster of similarities was again observed in delta-band power (P = 0.0270; Figure 2Bi), with neural activities between 750 and 1100 milliseconds after referent onset activated throughout the period of 0 to 1300ms post-pronoun onset, likely also extending to the pre-stimulus period.

Between-referent word control RSAs showed a significant activation of delta-band power patterns underlying referent word processing. The peak of the cluster arose between 650 and 950ms (P = 0.03; Figure 2Bii) and extended to later periods, sustaining until the end of the epoch period (i.e., 1.5 seconds). No significant effect was observed in the word control RSAs between pronouns. Source-level RSA analyses yielded no significant effect for neither the pronoun-referent or control comparisons (see Supplementary Materials for detail).

### 2 Theta band (4-7 Hz)

#### 1.1 Phase

In the pronoun-referent phase RSAs, a marginal but non-significant cluster where theta phase was more similar between mismatching referent and pronoun words than between matching word pairs was found (Figure 3Ai; P = 0.0508). Concretely, the cluster went between 200 and 550ms post-referent onset and predominantly between 350 and 700ms post-pronoun onset.

**Figure 3.**
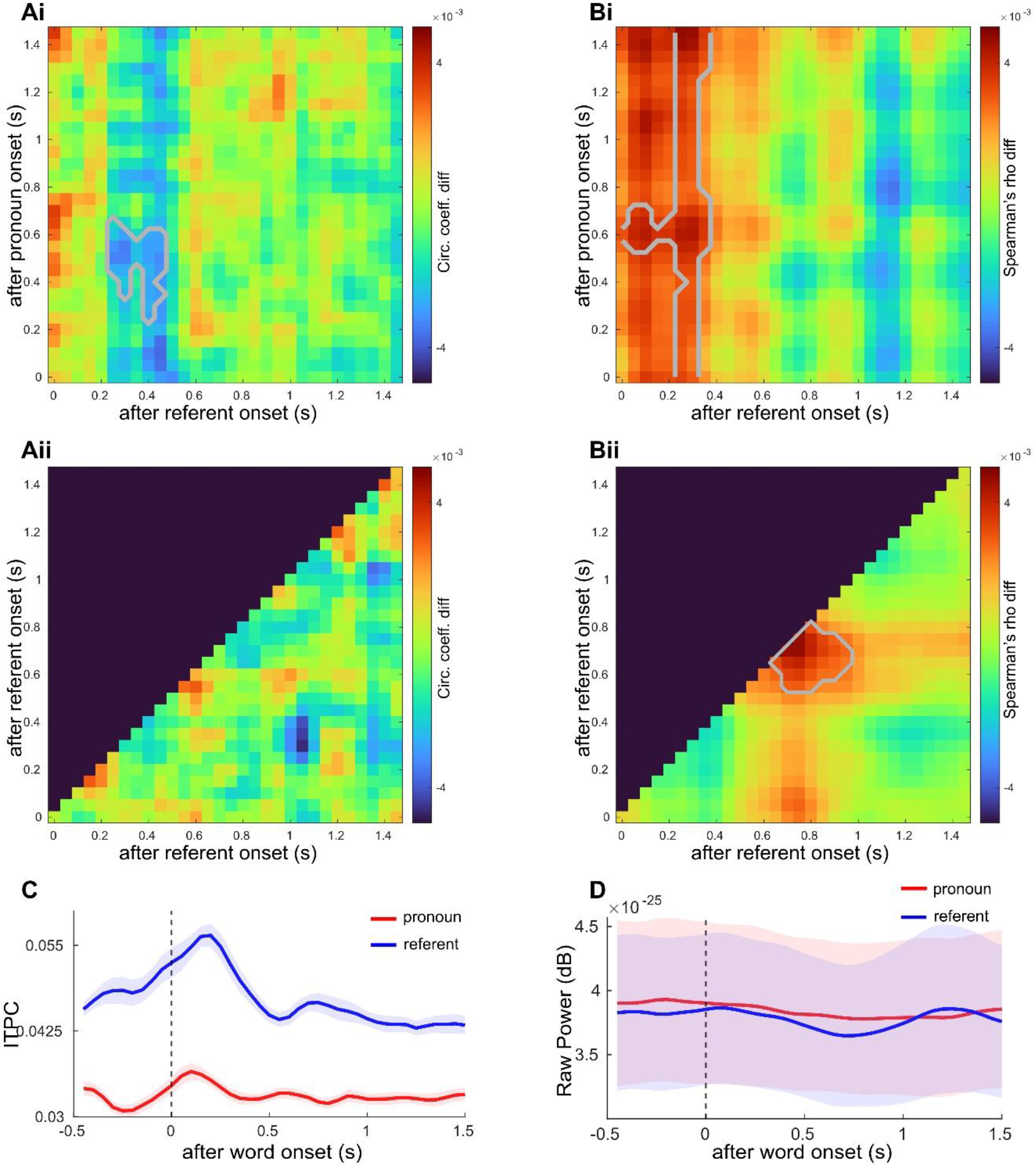
Reinstatement of theta-band (4-7Hz) activities during pronoun resolution. A) Temporal generalization maps averaged across participants for theta phase. (i) Averaged temporal generalization map for the difference between matching and non-matching pronoun-referent word pair conditions. (ii) Averaged temporal generalization map for the difference between formally identical and non-identical word pair conditions. B) Reinstatement of neural representations of referent words is found in theta-band power during pronoun resolution. (i) Averaged temporal generalization map for the difference between matching and non-matching pronoun-referent word pair conditions. (ii) Averaged temporal generalization map for the difference between formally identical and non-identical word pair conditions. C) Uncorrected, mean inter-trial phase coherence (ITPC) averaged per temporal channel per delta frequency. The ITPCs of pronoun and referent trials were plotted as separate lines. Note that the higher ITPC for referent words here could be due to the fact that the number of referent word trials are smaller than that of pronoun word trials. D) Mean delta power across the delta band and across all temporal channels. The mean power of pronoun and referent trials was plotted as separate lines. Shaded error bars indicate the standard error of the mean. Regions highlighted with gray outlines indicate trend-significant difference at the P = .05 level.

No significant effect was observed on theta phase in word control RSAs between referent words (Figure 3Aii) or between pronoun words.

#### 1.2 Power

The RSA between pronouns and referents yielded a trend-significance theta-band power reinstatement, primarily between 200 and 300 milliseconds after referent word onset and throughout the entire epoch (0-1500ms) of pronouns (P = 0.0543; Figure 3Bi). The cluster might extend to the pre-stimulus and post-epoch periods time-locked to pronoun onset.

Between-referent word control RSAs show a trend-significance pattern activation of theta-band power underlying referent word processing. The peak of the cluster arises between 600 and 950ms (P = 0.0942; Figure 3Bii). No significant effect is observed in the word control RSAs between pronouns.

### 3 Other frequency bands

No significant effect was found in any RSA in the alpha (8-12 Hz), beta (13-20 Hz), or low-gamma (35-80Hz) band.

## Discussion

When we understand a pronoun in a phrase, sentence, discourse, or story, the brain accesses matching entities or concepts that have been encoded in memory from the previously presented linguistic context. Recent advances in the neurobiology of memory point to a mechanistic account of memory retrieval wherein memory traces are (re)activated via the reinstatement of oscillatory neural representation underlying encoding (e.g., Rugg & Johnson, 2007). Meanwhile, in a cue-based account of language processing (e.g., Martin, 2016; Martin, 2020) it is proposed that, during pronoun resolution, the brain takes as its input exogenous cues (e.g., sensory features) and accesses via those external cues previously encoded, internal representations of entities that match the current pronoun, so that resolution of a pronoun is accomplished (LeDoux et al., 2007; Foraker & McElree, 2007; Parker, 2019). Importantly, in line with the findings of oscillatory dynamics reinstatement in memory retrieval, the account also asserts that internal linguistic elements can be represented by (a)synchronously firing neuronal ensembles. We therefore hypothesized that retrieving previously encoded antecedents engages the reinstatement of oscillatory neural dynamics as well during pronoun resolution. By leveraging state-of-the-art neural decoding techniques (i.e., representation similarity analysis) on MEG data acquired during a naturalistic story listening task, our study provides the first direct evidence of reinstatement of delta-band activity during pronoun resolution in ongoing spoken language comprehension. These findings suggest that the brain may retrieve a referent by reinstatement of its rhythmic neural representation, hereby serving as the first step to bridge the gap between neurobiological accounts of language processing and memory.

We assessed whether oscillatory activity underlying referent processing was reinstated during pronoun resolution. We took particular interest in delta-band (1-3Hz) activity as it has been shown to concern processing of higher-order, meaningful linguistic components such as words and phrases (e.g., Bai, Meyer, & Martin, 2022; Brennan & Martin, 2019; Coopmans et al., 2022; Keitel, Gross, & Kayser, 2018; Attaheri et al., 2022; Kaufeld et al., 2020; Lo et al., 2022; Meyer et al., 2017; Henk & Meyer, 2021; Molinaro & Lizarazu, 2018; Molinaro et al., 2016; Ten Oever et al., 2022; Weissbart, Kandylaki, & Reichenbach, 2020; see Meyer, 2017 for a review). Critically, we found strong effects of reinstatement over both the phase and power of antecedent-related delta-band responses during pronoun word processing, suggesting that delta-band dynamics may be involved as a mechanism that enables the brain to retrieve and represent an antecedent during language comprehension. Previous ERP results on the neural processing of reference have provided compelling supportive evidence for the cue-based account by showing that retrieving internal linguistic knowledge can be interfered by processing referentially incoherent or ambiguous input (e.g., Nieuwland & Van Berkum, 2008; Nieuwland, 2014; Karimi, Swaab, & Ferreira, 2018; Coopmans & Nieuwland, 2020). However, the current study takes an important first step further to characterise how such internal, abstract linguistic concept representations may be retrieved and involved in spoken language processing, namely via the reinstatement of delta range patterns. On this note, our results also add to prior evidence supporting the cue-based account and suggest that generation of linguistic elements is achieved via (a)synchronously firing of neuronal ensembles.

More crucially, our present results also speak to the emerging neurobiological account of memory processes in which neural patterns during encoding are reinstated at retrieval (e.g., Rugg & Johnson, 2007; Manning et al., 2012; Jang et al., 2017; Yaffe et al., 2017; Xiao et al., 2017; Pacheco Estefan et al., 2019; Staresina et al., 2019; Ten Oever et al., 2021). It has been proposed that, during encoding of an experience, both externally perceived and internally generated aspects of the experience are processed by sensory and association cortical areas (e.g., lateral and medial temporal lobes) and then integrated into a cohesive memory via the hippocampus (Mayes, Montaldi, & Migo, 2007; Preston & Eichenbaum, 2013). According to this proposal, to successfully access stored memory traces and re-experience the event, the brain has to reinstate relevant cortical activities (e.g., Preston & Eichenbaum, 2013; Sederberg et al., 2007; Jang et al., 2017). In support of the proposal, multiple recent studies have shown that the recall of an event involves reinstatement of oscillatory brain activity that occur during the encoding of the experience of the event (Yaffe et al., 2017; Staresina et al., 2019; Ten Oever et al., 2021). Pronoun resolution, in its essence, should be largely categorised as a memory retrieval process, as representations of a previously encoded antecedent would always have to be recovered based on sensory input of a pronoun. Therefore, by showing that neural-oscillatory patterns underlie the reinstatement of referent representations across temporal channels during pronoun resolution, our data converge with prior studies and suggest that retrieval of internally-stored linguistic knowledge also involves neural representation reinstatement. Note though that while our results are in line with prior findings that lower-frequency bands (e.g., theta) can be engaged in memory trace reinstatement, we did not observe any effect in the gamma range, which in contrast has been repeatedly found in the memory literature (e.g., Yaffe et al., 2014; Yaffe et al., 2017; Staresina et al., 2019; Ten Oever et al., 2021). This could have been due to the general low signal-to-noise ratio in gamma-band activity (Hansen et al., 2010) that makes it difficult to disentangle the gamma effect(s) from non-neural noise (e.g., muscle artifacts). It could also be the case that temporal alignment was not identical across trials, which had made correlations between gamma responses less distinguishable given the transiency of its cycles. Taken together, the current study moves one step forward to bridge theories of language and of memory in the brain that have long been studied separately. Future studies on language processing could further examine whether oscillatory neural (especially delta) pattern reinstatement also underlies other types of long-distance dependency resolution such as ellipsis or displacement arising from movement operations (e.g., *wh*-question construction in English), or even word composition in phrasal building.

Our findings also converge with the previous studies that show the involvement of delta-band activity in processing of higher-order, meaningful linguistic elements (e.g., Bai, Meyer, & Martin, 2022; Brennan & Martin, 2019; Coopmans et al., 2022; Ding et al., 2016; Henk & Meyer, 2021; Meyer et al., 2017; Ríos-López et al., 2020; Kaufeld et al., 2020; ten Oever et al., 2022; van Bree et al., 2021). However, earlier findings have not disentangled two potential processes that can involve the delta range, i.e. an intrinsic slower temporal constraint for the tracking of higher-order linguistic components from external speech streams (e.g., Rimmele et al., 2018; Lakatos, Gross, & Thut, 2019) or a functional pattern that facilitates the generation of abstract linguistic representations (e.g., Meyer, Sun, & Martin, 2019). Our results corroborate previous findings by showing that the delta frequency range is involved in processing of higher-level linguistic representations (i.e., referential meaning). More importantly, the current study goes one step further to show specifically that delta-band responses are engaged in the generation of top-down linguistic representations via reinstatement of information carried in phase and power, as referential meaning cannot be immediately “tracked” from the speech envelope but needs to be inferred based on internal knowledge and external input. We have to note, though, that our results do not rule out the possibility that delta activity is also involved in tracking higher-level linguistic input, as both phase and power patterns were also found between identical words at relatively earlier periods of processing than the time window of referential meaning retrieval. However, it is at least clear from the current study that the delta band does more than mere tracking, as recovering internally stored linguistic concepts is required for the brain to process a pronoun. Besides, differing patterns of reinstatement were identified in the present study between delta and theta, the frequency band that has been repeatedly found associated with syllable processing (e.g., Luo & Poeppel, 2007; Ten Oever & Sack, 2015; Kösem et al., 2018; Giraud & Poeppel, 2012; Poeppel & Assaneo, 2020) – namely that robust modulations of delta were observed during the resolution of a pronoun, but not of theta, even though pronouns are more often theta-sized than delta-sized. Concretely, we found that the time window of theta power reinstatement in referent words predominantly lasted from 200-350ms post-stimulus onset, whereas delta power reinstatement happened much later after referent word onset (i.e., 750-1100ms) (see Figures 2Bi and 3Bi). Such a distinction indicates that the delta frequency range tends to be engaged differently in spoken language processing than theta — that is, while theta may reflect early-period memory operations on speech streams such as syllable and word recognition, delta may be more closely involved in later processes, i.e. retrieval of more abstract linguistic representations (e.g., stored lexical information, knowledge of grammar, semantic memory, event representations, conceptual information). Our current findings provide empirical evidence to disentangle the role of delta activity in accessing internally stored linguistic representations from tracking external input during spoken language comprehension.

Zooming in to the current findings, intriguingly results of control analyses on antecedent words showed a consistent temporal relationship with the pronoun-referent results in both delta phase and power. For delta phase, the reinstatement effect, which likely suggests referential meaning retrieval, was identified between 350 and 700 milliseconds after referent onset. Meanwhile, control RSAs between antecedent words yielded an earlier, trend-significance delta-band phase pattern activation between 200 and 450ms time-locked to antecedent word onset, which interestingly is consistent with the time window of the N400 ERP effect which has been suggested to be associated with processing at the lexical level (e.g., lexico-semantic, phonological; Kutas & Hillyard, 1984; Van Berkum et al., 2005; DeLong, Urbach, & Kutas, 2005; Frank et al., 2015; Ito et al., 2016). Therefore, this indicates that, for referent words, (pre-)lexical processing takes place earlier than referential meaning retrieval in referents. This is convergent with prior findings suggesting a sequential model that referential resolution follows lexical activation (e.g., Brodbeck, Gwilliam, & Pylkkanen, 2016; Coopmans & Nieuwland, 2020). For delta power, again the onset of the pattern activation induced by formally identical referent words (i.e., starting from 600ms after referent onset) was found to precede that of the reinstatement effect identified in referents (i.e., 750 and 1100 milliseconds after referent onset). Note, though, that in delta power the effect of reinstatement ran throughout the entire epoch period after pronoun onset, likely also extending to the pre-stimulus period (see Figure 2Bi). We still regard it as reinstatement given that the pre-stimulus similarities we observed in pronouns are unlikely to reflect activities that follow from those induced by an immediately preceding referent word (we excluded all datapoints where the time between two words in a pair was shorter than 1.5 seconds). Such a sustained effect in pronouns may suggest that referential representation stays retained in working memory when the brain processes ongoing speech. It is also interesting to note that, while the inter-trial phase coherence (ITPC) of delta activity was clearly distinct between pronoun and referent trials, the mean power seemed to stay highly similar (see Figure 2C and D). The higher ITPC in referents than pronouns could be due to the fact that referent trials are smaller in number. But the discrepancy between ITPC and power indicates that phase measures can be more sensitive to conditional contrasts (e.g., adjusts more transiently to sensory input) than power in naturalistic experimental setups. Taken together, these results provide compelling evidence that delta-band activity is involved in late, post-lexical construction of referential meaning during pronoun resolution, more importantly via reinstatement. This is also consistent with the cue-based account of language processing which claims that internally stored linguistic cues guide linguistic structure generation in slower neural responses such as delta (Martin, 2020).

Note that we did not found any significant effect in our RSA analysis on the source level (see Supplement 1 for method). Specifically, we performed ROI-based RSA in the delta band to test whether dipoles in the temporal areas were significantly involved in the reinstatement effect that we observed in the sensor space. No effect was found in either lateral or medial temporal region (see Supplement 2), indicating that the reinstatement effects we identified on the sensor level did not take place at least only in temporal areas. Indeed, our control analyses across all channels in the sensor space showed that the reinstatement effect on power were stronger and longer-lasting after referent word onset (see Supplement 3). This suggests that the delta-band activity involved in reinstatement of referential meaning may be distributed across the cortex.

Yet, it is still worth pointing out that the exact functional role delta activity plays in neural representation of reference remains an open question. Pronoun resolution is inherently difficult to separate from other cognitive processes associated with building or updating the situation model of a discourse, which includes for instance integration of referential entities to the event where they are embedded, or encoding of new features into entity representation. Therefore, it is possible that the observed delta-band reinstatement does not precisely reflect the representation of any individual referential concept in isolation, but a more dynamic process that involves relational concept representation. One potential alternative interpretation is that delta reinstatement reflects a referential entity-specific binding process where the entity is integrated with its predicate or broader sentential context as an argument (as in Martin & Doumas, 2017 for example, see Martin & Doumas, 2019 for a review). Another way to look at the delta reinstatement effect is that previously encoded features of an entity may be synchronously replayed in delta cycles so that the features are subject to interference by incoming novel information. Although there is as of yet little empirical evidence showing the involvement of delta activity in memory replay during the awakening state (see e.g. Girardeau & Lopes-Dos-Santos, 2021 for a review on delta replay during sleep), slower oscillations such as theta in the hippocampus have long been found to be associated with episodic memory formation and consolidation (e.g., Jensen & Lisman, 2005, 2013). Meanwhile, it has also been suggested that hippocampal theta oscillations tend to be slower in humans, i.e., fall into 1-4Hz, the frequency range usually named as delta (e.g., Jacobs, 2014). It is therefore reasonable to speculate that delta activity could also be engaged in memory formation. Our data cannot distinguish between these possibilities. We invite future research to investigate the functional role of the delta frequency range in the process of pronoun resolution more closely.

In summary, by bridging bourgeoning, brain oscillation-based accounts of language processing and memory, our findings provide new insight into the neurobiological substrates of referent representation and pronoun resolution. We show that establishing reference involves the reinstatement of neural patterns that first occurred during the processing of the antecedent word, in particular occurring in the delta band. Our data thus suggest that, during spoken language comprehension, processing that calls upon internally-stored linguistic representations in order to create meanings (here, referential dependencies) may require some degree of reinstatement of the oscillatory brain responses that occurred when those stored representations first encountered.

