## Supplement 1 for "Pronoun resolution via reinstatement of referent-related activity in the delta band"

Supplement 1 MEG source reconstruction & Source-level RSAs: Method

*ROI-based RSAs*. Source-level analyses were conducted to investigate the cortical source(s) of the reinstatement effect identified in the sensor space. To examine the hypotheses that the temporal lobe is engaged in the reinstatement of neural activities underlying referent representation, we performed ROI-based RSAs in the delta band on the source level to test whether dipoles in the temporal areas were significantly involved in the reinstatement effect that we observed in the sensor space. The linearly constrained maximum variance (LCMV) beamformer approach (Van Veen et al., 1997) was used to obtain the source reconstruction of each single trial. A spatial normalisation with the array-gain method was applied to mitigate the centre-of-head bias (Sekihara & Nagarajan, 2008; Westner et al., 2022). The data covariance matrix was computed over each entire trial epoch (-0.5 to 2 seconds time-locked to stimulus onset), and was subsequently used, in combination with a forward model defined on a set of source locations (8004 in total), to construct common spatial filters, each of which corresponds to one dipole location. Channel-level trial data were then projected to the source space through the spatial filters. Individual cortical sheets and dipole parcellation were produced with the Freesurfer package (Dale et al., 1999; version 7.2.0). The forward model was generated using *singleshell* method in Fieldtrip (Nolte, 2003), where the skull/brain boundary information was extracted from the T1-weighted MRI scan of each participant. For the lateral temporal ROI, we selected on the basis of individual cortical sheets the parcels that form bilateral lateral temporal areas (i.e., superior, middle and inferior temporal gyri, fusiform gyri, transversetemporal gyri, temporal pole, bank of superior temporal sulci) as components of the ROI.

Note that, to estimate the source power of medial temporal areas (which was not supported by the cortical sheet-based method), we additionally implemented a volume conduction source modelling method. In volume conduction modelling, the forward model with a total of 11000 source locations was generated by warping a template MNI grid to individual resliced MRI scan that had been transformed to the ctf-format in Fieldtrip. To generate tissue labels for each dipole location, a template atlas was interpolated upon the template MNI grid. Then, for the medial temporal ROI, tissue labels that comprise the bilateral medial temporal regions (i.e., hippocampus, parahippocampal area, and amygdala) were selected as components of the ROI. The same LCMV beamformer approach as delineated above was conducted after a volume conduction model was created.

After source estimation was completed, time-frequency analyses on were conducted on the trial data of each dipole within the ROIs. Again, the activities in source dipoles underwent a wavelet analysis whose procedure was identical to what had been conducted on the sensor level. The time-frequency results (phase or power) were then subjected to a sliding-window RSA, and then the similarity values of either condition (matching or non-matching) in the time*time points in the statistically significant cluster that had been identified in the corresponding sensor-level analysis were selected and averaged per participant respectively. Subsequently, a dependent-sample t-test was conducted for the phase or power of each ROI on the averaged matching and non-matching datapoints across participants.
