## Supplement 2 for "Pronoun resolution via reinstatement of referent-related activity in the delta band"

Supplement 2 Source-level ROI-based RSAs: Results

As described in Supplement 2, RSAs were conducted on the lateral and temporal ROIs separately. On each ROI we performed the analysis on delta-band phase and power respectively.

Dependent-samples t-tests on the RSAs based on the lateral temporal ROI suggest no significant difference between matching and non-matching conditions in either delta phase ($t= 0.505$, $p=0.6194$) or power ($t=-1.3117$, $p=0.2015$) in lateral temporal regions. No significant difference is found in medial temporal regions either between matching and non-matching conditions in delta phase ($t=-0.6282$, $p=0.5356$) or power ($t= -0.885$, $p=0.3844$).
