## Supplement 3 for "Pronoun resolution via reinstatement of referent-related activity in the delta band"

### Supplement 3 Sensor-level RSA results in the delta band across all channels

From the RSA between pronouns and referents, a significant cluster of similarities was observed in delta-band power ($P = 0.012$; Supplementary Figure 1A). Word control RSAs between referent words yielded a significant delta-band power pattern activation (Supplementary Figure 1B; $P = 0.0225$). The control RSA between pronoun words showed a trend-significant pattern activation cluster (Supplementary Figure 1C; $P = 0.0519$) and a significant pattern de-activation cluster ($P = 0.0478$).

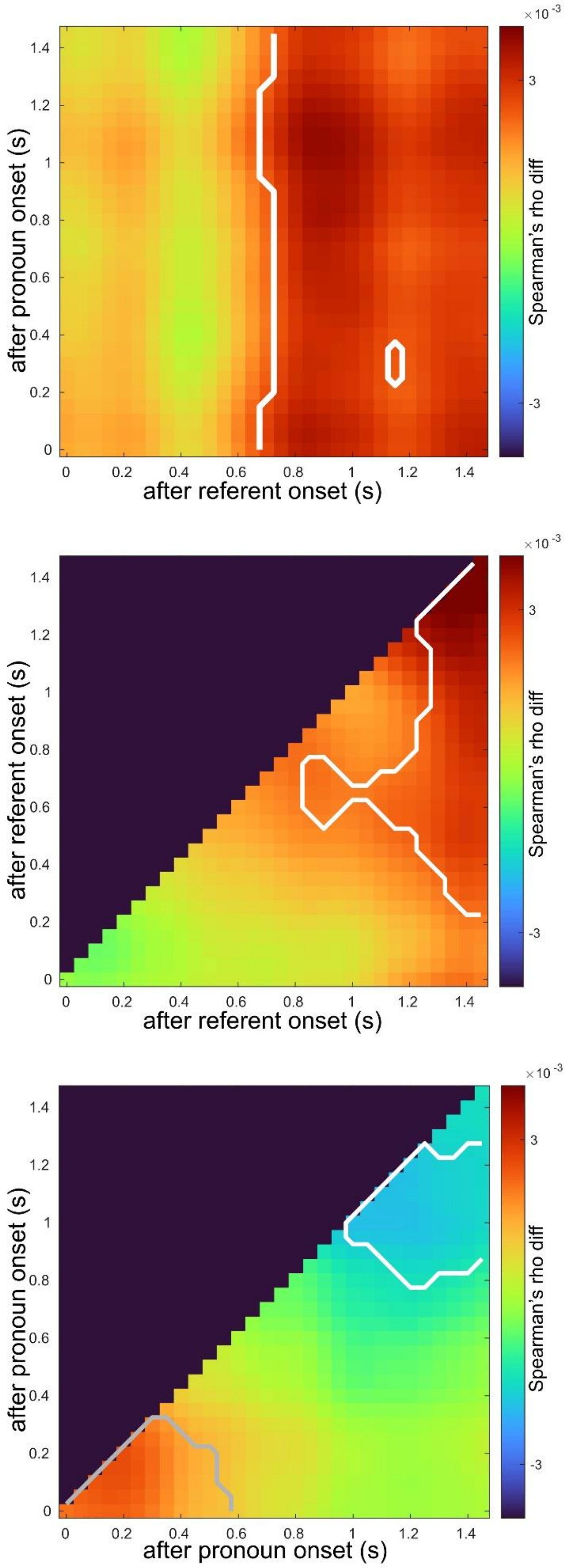

after pronoun onset (s)
after referent onset (s)
Spearman's rho diff
×10^-3
3
-3
0
0.2
0.4
0.6
0.8
1
1.2
1.4
after referent onset (s)
after referent onset (s)
Spearman's rho diff
×10^-3
after pronoun onset (s)
after pronoun onset (s)
Spearman's rho diff
×10^-3

**Supplementary Figure 1 Reinstatement of delta-band (1-3Hz) power during pronoun resolution across all MEG channels**.

A) Temporal generalization map averaged across participants for the difference between matching and non-matching pronoun-referent word pair conditions. Regions highlighted with white outlines indicate significant difference at the P = .05 level, while those highlighted in gray indicate trend-significant difference.

B) Temporal generalization map for the difference between formally identical and non-identical referent word pair conditions.

C) Temporal generalization map for the difference between formally identical and non-identical pronoun word pair conditions.
